# Shuffle-Seq: *En masse* combinatorial encoding for *n*-way genetic interaction screens

**DOI:** 10.1101/861443

**Authors:** Atray Dixit, Olena Kuksenko, David Feldman, Aviv Regev

## Abstract

Genetic interactions, defined as the non-additive phenotypic impact of combinations of genes, are a hallmark of the mapping from genotype to phenotype. However, genetic interactions remain challenging to systematically test given the massive number of possible combinations. In particular, while large-scale screening efforts in yeast have quantified pairwise interactions that affect cell viability, or synthetic lethality, between all pairs of genes as well as for a limited number of three-way interactions, it has previously been intractable to perform the large screens needed to comprehensively assess interactions in a mammalian genome. Here, we develop Shuffle-Seq, a scalable method to assay genetic interactions. Shuffle-Seq leverages the co-inheritance of genetically encoded barcodes in dividing cells and can scale in proportion to sequencing throughput. We demonstrate the technical validity of Shuffle-Seq and apply it to screening for mechanisms underlying drug resistance in a melanoma model. Shuffle-Seq should allow screens of hundreds of millions of combinatorial perturbations and facilitate the understanding of genetic dependencies and drug sensitivities.

---

Genetic interactions occur when the combined impact of *n* genes cannot be determined by their additive individual impacts. A comprehensive test for interactions requires generating datasets of a sufficient scale to test for all possible interactions between the genes within a sizeable set, with the number of combinations growing as 2^n^ with *n* being the number of genes in the set or _n_C_2_ for pairwise interactions only. A common example phenotype is cell fitness, where an extreme case of a genetic interaction is synthetic lethality. In yeast, all pairs of genetic deletions have been constructed over the course of a decade and quantified for their relative fitness impact^1–3^, with recent studies targeting a small portion of all three gene deletions^4^. While it is likely that the total number of significant interactions, with respect to fitness in yeast, is far larger numerically for three-way interactions than pairwise, their prevalence is likely lower (3% of pairs as opposed to 1% for three-way). Their scarcity does not preclude their importance; these higher-order genetic interactions can play a major role in the evolution of genetic networks^5,6^, and have been translationally leveraged for cellular reprogramming^7^. In mammalian systems, studies have examined the impact of pairwise gene interactions only for small subsets of genes (between 25 and 323), assessing cell growth^8–11^, drug resistance^10,12^, cell morphology^13^, or gene expression^14–17^.

Because the potential number of combinations to screen is vast (>10^8^ and >10^12^ for all 2- and 3-way combinations of genes in the human genome) it has been challenging to perform comprehensive screens. Broadly, existing methods either (**1**) create each interaction independently in massive arrays (*e.g.*, in yeast^1–3^), (**2**) create a single barcoded vector containing multiple perturbations^8–11,27^, or (**3**) use single cell profiling to determine, *post hoc*, the set of perturbations present in a cell^14–17^. They are respectively limited by (**1**) time, reagent, and labor costs required to scale, (**2**) scale and length limitations on oligo synthesis, titer decreases associated with longer insert lengths^18^, decreased stability as more perturbations are introduced^19^, and the inability to simultaneously test interactions between orthogonal types of perturbations like overexpression and knockout libraries, and (**3**) the cost of scRNA-seq and similar methods^14^ (**Supplementary Table 1**).

To tackle these limitations for mammalian screens – including viability and FACS-based screens – we developed Shuffle-Seq, combining the benefits of coupling perturbations at a single cell level with the scale of pooled screens (**Fig. 1a,b**). Shuffle-Seq uses Unique Transduction Barcodes (UTBs), similar to those used in previous CRISPR screens^20,21^, such that the probability of two cells getting the same combination of perturbation-UTBs is extremely low. Multiple perturbations are delivered into the cell through one or more rounds of viral transduction. The cells are allowed to expand in a pool and subsets from the pool of cells from the multiple resulting clonal populations are randomly distributed into standard multiwell plates. Barcoded perturbation pairs that were delivered to the same transduced cell, will propagate to its progeny prior to the distribution step, and, as a result, their probability of co-occurrence across wells is significantly higher than for random pairs (not from the same cell) (**Supplementary Fig. 2b,e,f**). By co-association across wells, pairs can be merged together to reconstruct the set of perturbations present within each clone. Thus, we can infer the clonal origin of sets of perturbations from the patterns of their co-occurrence across wells (**Fig. 1a, bottom**). This inference, for *n* barcodes, is analogous to an approach developed for pairing endogenously expressed T cell receptors^22^. Finally, we identify, as in any screen, combinations of perturbations that are enriched or depleted. Shuffle-seq can be applied after a strong positive selection, such as FACS based gating, or in a negative selection screen, where it is performed shortly after the introduction of perturbations to establish clonal pairings, followed by bulk sequencing to observe how clonal frequencies change over time (**Fig. 1b**).

**Figure 1.**
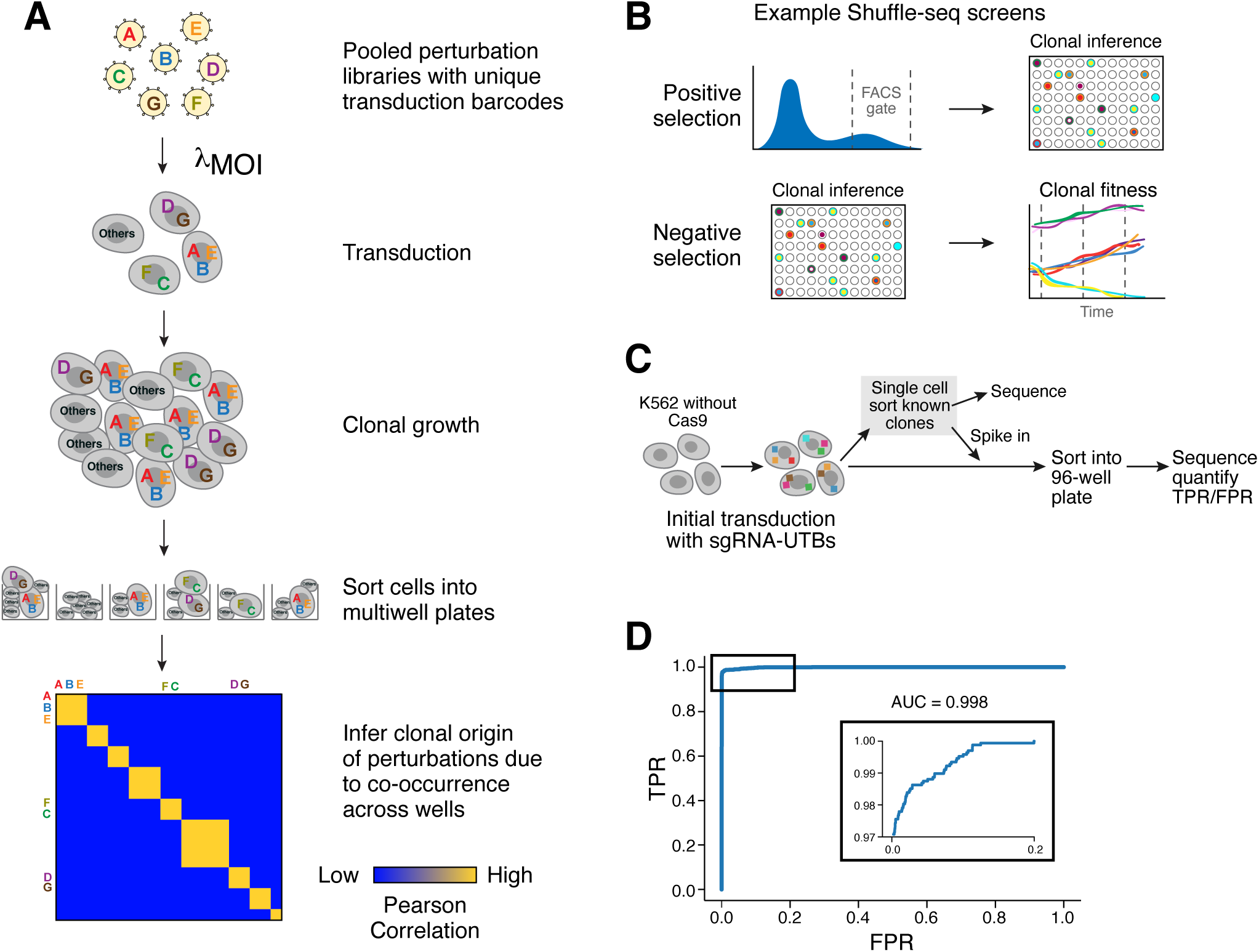
Shuffle Seq approach. (**a**) Shuffle-seq. Cells are transduced with virus (at high MOI and/or multiple rounds with orthogonal selectable markers), such that most have more than one perturbation, and allowed to clonally expand. After a screening assay, cells are sorted into a multiwell plate with multiple cells per well (10 to >10,000 cells/well). Wells are profiled by targeted amplicon sequencing of unique perturbation barcodes. Clonal origin is inferred from correlated patterns of perturbation identities across wells, provided the library size is significantly greater than the number of clones to be inferred. (**b**) Shuffle-seq can be applied in multiple screening contexts, including positive and negative selection screens. Top: in positive selection, Shuffle-Seq is performed (right) after sorting of positive cells (left). Bottom: In negative selection, Shuffle-Seq is performed first (left), followed by bulk targeted amplicon sequencing of the unique perturbation barcodes over time (right). (**c,d**) Clone inference with Shuffle-Seq. (**c**) Experimental design. K562 cells without Cas9 were transduced by an sgRNA Shuffle-Seq library. 88 clones were sorted, grown, and representative cells were either profiled (top path) or spiked-in back to the pool at different concentrations, prior to Shuffle-Seq (bottom path). (**d**) Receiver operating characteristic (ROC) curve of the True Positive Rate (TPR) (*y* axis; proportion of sgRNA-UTB pairs inferred by Shuffle-seq out of the pairs ascertained in the 88 positive controls) and False Positive Rate (FPR) (*x* axis; proportion of pairs from distinct clones that were co-associated across Shuffle-seq wells) (**Methods**). Inset: Zoom in to the region between FPR 0.0 and 0.2.

**Figure 2.**
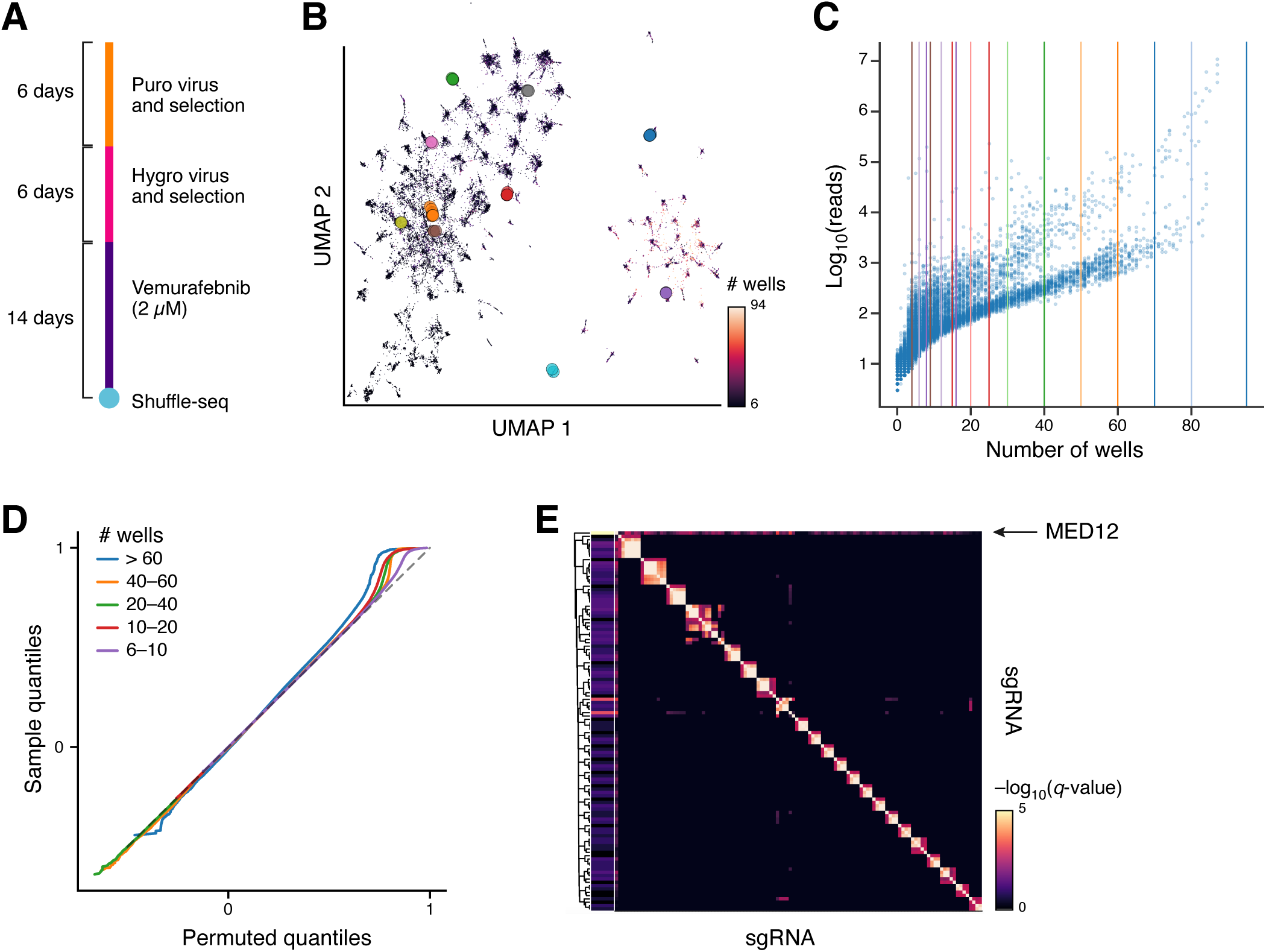
Shuffle-seq in a Vemurafenib resistance screen in a melanoma cell line. (**a**) Screen overview. (**b-d**) Clone inference (**Methods**). (b) Similarity structure of sgRNA-UTB abundances across wells. UMAP embedding of sgRNA-UTBs (dots) occurrence profiles across wells colored by number of wells in which the sgRNA-UTB is detected. Larger dots correspond to sgRNA-UTBs from ten randomly selected clones, colored by clone. (c) sgRNA-UTB distribution across wells. Number of wells (y axis) and number of reads (log_10_(total number of reads), *x* axis) for each sgRNA-UTBs. (d) Enrichment in pairwise correlation of sgRNA-UTB pairs relative to permuted sgRNA-UTB pairs. Q-Q plot of the Pearson correlation coefficient between pairs of sgRNA-UTBs of various abundances in the real data (*y* axis) and in a permuted sgRNA-UTB abundance matrix (*x* axis). (**e**) sgRNA interaction matrix of significant co-occurrences across clones. Significance (log_10_(hypergeometric q-value), color bar) of co-occurrence for sgRNA pairs (rows, columns). sgRNAs have been subset to the those with at least four significant interactions. Left colorbar: log_10_(individual read abundance) of each sgRNA in the Shuffle-seq library.

To maximize the detection of the perturbation-UTB, we modified the CROP-seq plasmid^17^, which generates a polyadenylated sgRNA transcript, to include an optimized trRNA^23^ and a UTB (**Supplementary Fig. 1a**). These modifications are introduced during the sgRNA cloning step (**Supplementary Fig. 1b**) and do not reduce cutting efficiency (**Supplementary Fig. 1c**). Sequencing of the sgRNA-UTB library shows that the number of distinct sgRNA-UTB pairs likely exceeds 50 million (**Supplementary Fig. 1d,e**).

We optimized experimental and computational approaches for clonal inference. If clonal abundances are biased, as may happen due to selection, an over-sampled clone might be present in all wells, while an under-sampled clone would be present in few or no wells. To mitigate this, we devised a modified procedure, akin to high dynamic range (HDR) photography, in which different numbers of cells are sorted per well – such that some wells have many more cells than others – in order to represent each clone in an intermediate number of wells (**Supplementary Fig. 2b-d**). Bulk sequencing of the perturbation library in advance of the Shuffle-seq experiment can inform the sort procedure. We performed simulations to determine the relationship between our statistical power to infer clones and the sgRNA-UTB detection probability and number of unique barcodes. We found that for reasonable detection efficiency and sequencing depth, a 96 well plate can be used to infer over a million clonally shared barcodes, while a 384 well plate can be used to infer over a billion (**Supplementary Fig. 2e,f, Methods**).

We next validated that Shuffle-seq can infer clones independently of fitness effects associated with the perturbations. We transduced our high complexity sgRNAs-UTB lentiviral library into K562 cells without Cas9, and, as positive controls, isolated 88 single cell clones from the transduced pool, and sequenced their sgRNAs-UTBs. In parallel, we reintroduced cells from these 88 clones to the parental pool at varying concentrations, and then performed Shuffle-seq on the entire pool (**Fig. 1c**). Based on the positive control clones, we obtained an AUC of 0.998 for recovering known pairs of sgRNA-UTBs from a background of pairs derived from a randomly permuted set of sgRNA-UTB abundances across the ∼300,000 other clones present in the wells (**Fig. 1d**). Our results were robust to the number of perturbations per cell, the expression level of the sgRNAs-UTB, and the number of wells a clone was present in (**Supplementary Fig. 3c,d**). We estimated our probability of detecting an sgRNA-UTB at ∼78% per well, based on the percentage of wells with co-occurrence of sgRNA-UTBs that were ascertained as co-occuring in one of the 88 clones (**Supplementary Fig. 3e**).

To show the feasibility of applying Shuffle-Seq in a biological perturbation experiment, we used the model system of resistance of the melanoma A375 cell line to the B-Raf inhibitor vemurafenib (**Fig. 2a**), which had previously been used for loss-of-function screens^24^ We infected cells in two rounds (to ensure at least two perturbations per cell) with a genome wide lentiviral library of 51,467 sgRNAs depleted for essential genes and including 5,000 non-targeting and 5,000 intergenic sgRNAs. In each round, we combined pools of cells transduced at viral MOI of approximately 1, 6, and 16 (to obtain a wide distribution of perturbations per clone). The first round used a vector containing puromycin resistance followed by a second round using a vector containing hygromycin resistance. During the two rounds selection, the cells were bottlenecked to approximately 100,000 clones. After positive selection with vemurafenib, we performed Shuffle-seq, in HDR mode, sorting between 25 and 20,000 cells per well.

We inferred 1,765 sgRNA-UTB pairs with shared clonal origin (**Fig. 2b-d**), even though the virus library constructed was bottlenecked relative to the initial plasmid pool and our culture scale was small relative to the library size resulting in poor representation of different genetic perturbations in our screen (**Supplementary Fig. 4a**). The inferred pairs were enriched for genes that are highly expressed in A375 cells (p=8.6*10^−39^) (**Supplementary Fig. 4e**). Grouping pairs that shared an sgRNA-UTBs, we consolidated to 597 inferred clones with 2-9 sgRNA-UTBs per cell (**Supplementary Fig. 4b**). Most (92%) of these inferred clones contained an sgRNA targeting MED12, a hit from previous CRISPR screens of vemurafenib resistance. Another 432 pairs of sgRNAs that did not contain MED12 were significantly enriched for co-occurrence across the clones (hypergeometric test, q-val <0.05, **Fig. 2e**).

Shuffle-seq enables large scale combinatorial screening by leveraging the encoding capacity of multiwell plates and the throughput of sequencing. In a proof-of-concept demonstration, it recovered clones, associating multiple barcodes, and generated hypotheses about potential new interactions that may underlie resistance in a melanoma cell lines; these interactions can be tested experimentally. In contrast to traditional approaches that operate in a regime where each cell contains a single distinct perturbation, Shuffle-Seq opens the way to high-throughput combinatorial screening in which each cell contains larger numbers of perturbations. While a large number of double strand breaks from multiple CRISPR/Cas9 perturbations could cause toxicity^25^, Shuffle-Seq can be generally applied to any perturbations that can stably propagate alongside their affected cell, including CRISPRi, overexpression, and variant ORF libraries. Uniquely, the approach can couple combinations of these perturbations from distinct vector constructs. The clonal inference in Shuffle-seq can also be extended to enhance detection in pooled high dimensional single cell screens in primary cells, where conventional detection of the perturbation in an individual cell can be lower than 60%^14^, to allow more accurate reconstruction of the initial set of perturbations present in cells sharing an sgRNA-UTB.

For those phenotypes, like drug resistance, where we expected only a small number of perturbations to have an impact, Shuffle-seq may be applied in the context of group testing and compressed sensing of perturbations^26^ to rapidly identify the relevant synergies (**Supplementary Fig. 2g**). We expect sequencing costs to be the main limiting factor associated with the total number of clonally associated perturbations that can be inferred with a Shuffle-seq experiment. As such, Shuffle-seq has the potential to facilitate the screening of hundreds of millions of clones, each of which can contain multiple perturbations, and should accelerate the discovery of new synergistic combinations that lead to translationally relevant phenotypes.

## Acknowledgements

We thank Brian Cleary for brainstorming the spike-in proof of concept experiment in K562 cells. Work was supported by the Klarman Cell Observatory, Howard Hughes Medical Institute, and a gift from the Wertheimer Family Foundation. A.R. is a founder of and equity holder in Celsius Therapeutics, an equity holder in Immunitas, and an SAB member of ThermoFisher Scientific, Syros Pharmaceutical, Asimov, and Neogene Therapeutics. A.D. is a founder of and equity holder in Coral Genomics. A.D. and A.R. have filed a patent application related to Shuffle-seq.

## Author Contributions

A.D. and D.F. conceived of the Shuffle-seq approach. O.K. performed the experiments with guidance from A.D. and A.R. A.D. performed the analysis. A.D., O.K, and A.R. wrote the manuscript with input from D.F.

## Supplementary Figure Legends

**Supplementary Figure 1.**
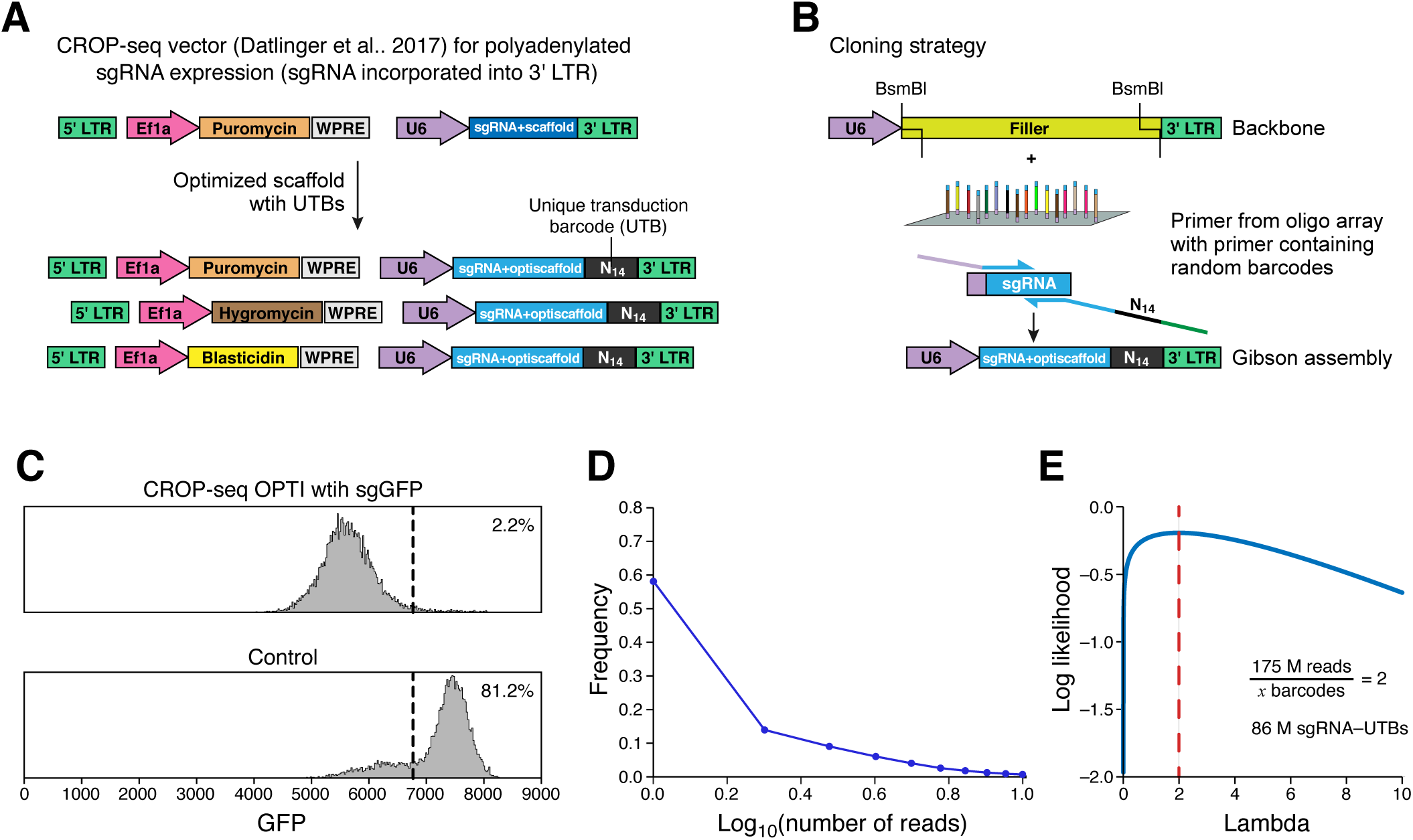
The Shuffle-Seq vector. (**a**) Shuffle-Seq vector. The CROP-seq vector^17^ (top) was modified to include an optimized scaffold^23^, a 3’ 14 bp random barcode to serve as the UTB, and one of several alternative selection markers (bottom). (**b**) Cloning strategy. The scaffold and barcode are added onto an oligonucleotide array of an sgRNA library during a PCR stem. (**c**) Successful knockdown by the modified vector. Distribution of GFP levels (*x* axis) in Cas9-P2A-GFP K562 cells transduced with the modified (top) or control (bottom) vector, showing 97% reduction in GFP expression. (**d**) High detection of the Shuffle-Seq sgRNA-UTB. Distribution of number of reads per sgRNA-UTB pair (*x* axis, log_10_(number of reads)) after deep sequencing of the high complexity sgRNA-UTB library. (**e**) Estimation of Shuffle-Seq sgRNA-UTB complexity. Log-likelihood (*y* axis) fit of the observed distribution of reads to a Poisson distribution (*λ*, y axis). Bottom right: calculation to infer that at least 86 million barcodes are present in the CRISPR-KO library.

**Supplementary Figure 2.**
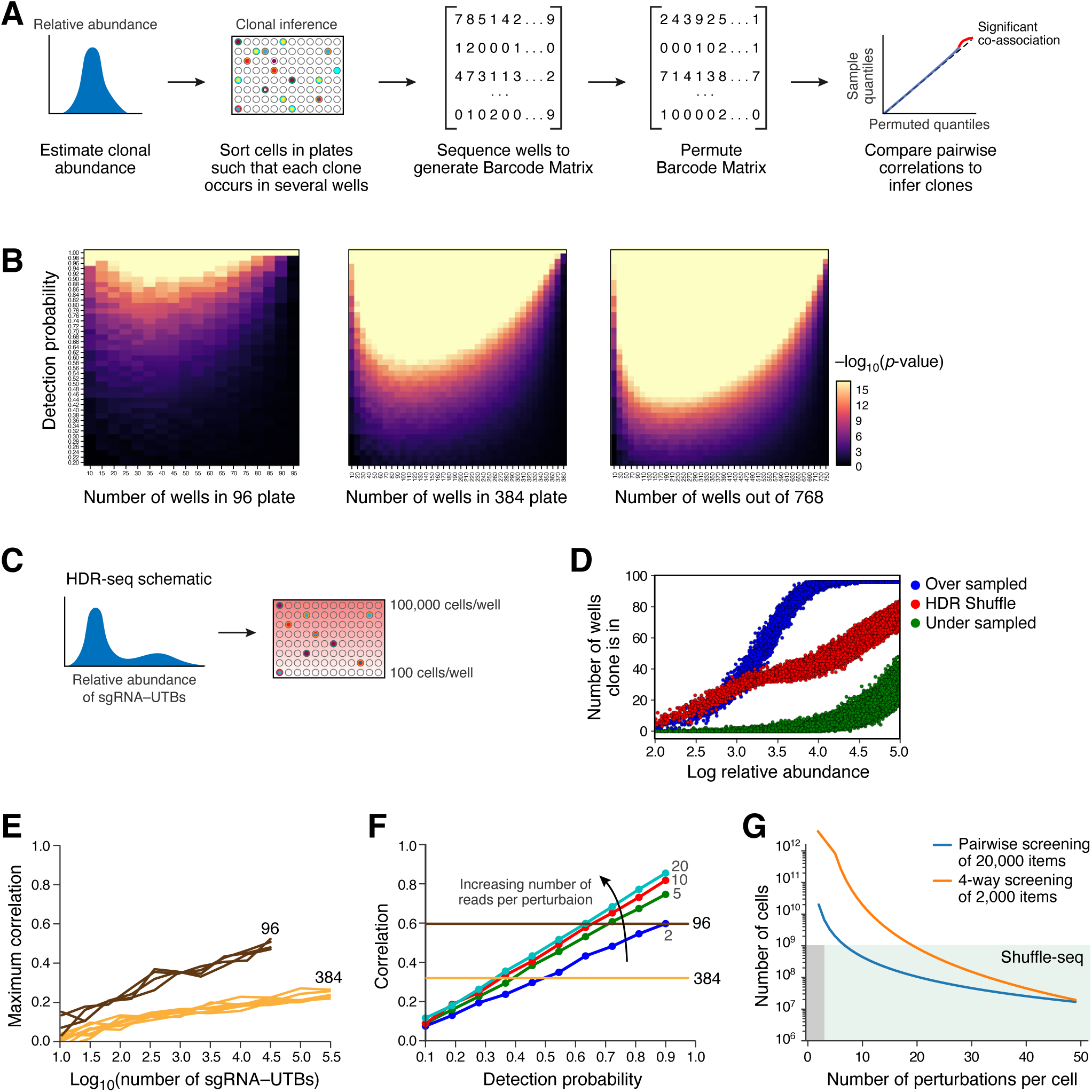
Simulation based assessment of Shuffle-seq approach. (**a**) Overview of approach. From right: Relative abundance of each sgRNA-UTB in the library overall is estimated based on bulk sequencing. Cells are sorted into a multiwell plate, such that each clone is expected to be represented in at least 5 wells. Reverse transcription and PCR are used to identify which sgRNA-UTBs occur in which wells. Pair-wise correlations in well occurrence profiles are calculated for each pair of sgRNA-UTBs, to identify significantly co-occurring barcode pairs (based on a permutation test associated with a randomly permuted matrix of sgRNA-UTBs across wells) and infer clonal identity. (**b**) The number of wells a clone must be represented in for accurate detection depends on number of wells and detection probability. Significance (-log_10_(P-value), hypergeometric test, color bar) for the co-occurrence of a pair of sgRNA-UTB from the same clone for different probabilities of sgRNA-UTB detection (*y* axis) when the clone is present at varying number of wells (*x* axis) out of 96 (left), 384 (middle) and 768 (right) wells overall. (**c,d**) HDR sorting strategy can address skewed sgRNA-UTB abundances. (**c**) Illustrative sort strategy. (**d**) Number of wells in which each clone is present (*y* axis) for clones (dots) of different abundance (*x* axis) in a highly skewed pool of clones sorted uniformly such that rare clones (green) are under sampled and very abundant clones (blue) are oversampled, vs. in HDR Shuffle-Seq (red). (**e**) Incorporation of sgRNA-UTB abundance per well. The maximum correlation (*y* axis) between two Poisson distributed vectors across a 96 (purple) or 384 (orange) well plate for different total number of sgRNA-UTBs present (*x* axis). (**f**) The correlation coefficient in quantitative abundances (*y* axis) between two sgRNA-UTBs across wells derived from the same cells at different detection probabilities (*x* axis) and sequencing depths (colors). (**g**) Representation advantage of testing large sets of perturbations when higher order interactions are rare. Approximate number of cells required to represent all items of a screen (*y* axis, 100x the total number of parameters) for different numbers of perturbations in each cell (*x* axis). Dark grey shaded area: traditional screens. Light grey area: screens enabled by Shuffle-seq.

**Supplementary Figure 3.**
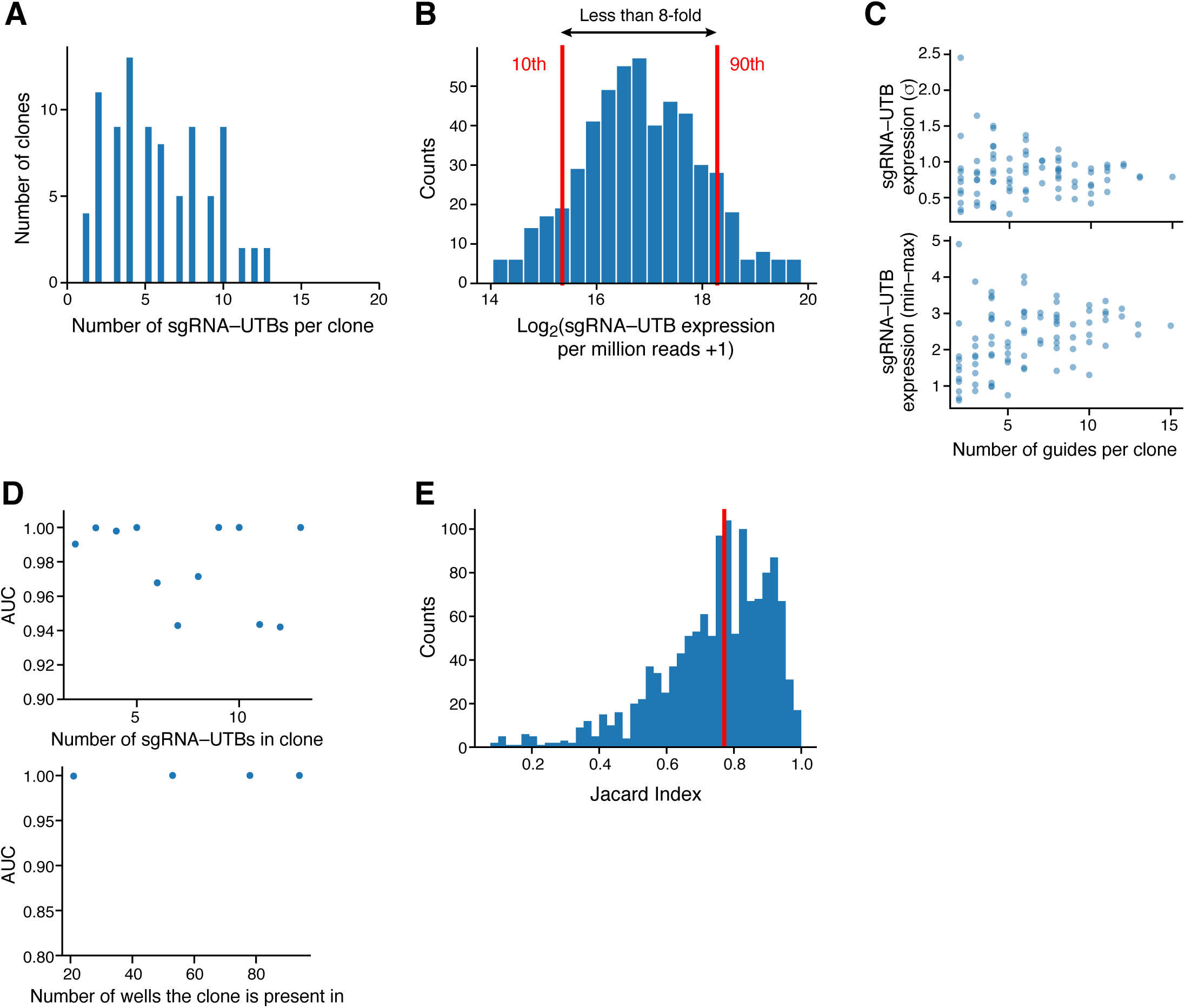
Statistics of a K562 Shuffle-Seq assay to infer clones without gene perturbation. (**a-c**) Positive controls. (**a**) Distribution of the number of sgRNA-UTBs observed in each of 88 derived single cell clones. (**b**) Expression of sgRNA-UTB. Distribution of expression levels (log_2_(TPM+1), *x* axis) of sgRNA-UTBs across 88 clones. (**c**) The number of sgRNA-UTBs in the clone (*x* axis) is not significantly correlated (r=-0.14, p=0.19) to the standard deviation of sgRNA-UTB expression (*y* axis, top) and is at most weakly correlated (r=0.19, p=0.07) to the range of expression (max – min, *y* axis, bottom). (**d**) Accuracy of clone detection does not strongly depend on the number of sgRNA-UTBs per clone and the number of wells the clone is present in. Area Under the Curve (AUC) (*y* a*x*is) – associated with **Fig. 1d** - for clones with different numbers of sgRNA-UTBs (*x* axis, top) or present in different numbers of wells (*x* axis, bottom). (**e**) Sensitivity of clone detection. Distribution of Jaccard overlap across wells (*x* axis) between sgRNA-UTBs belonging to the same clone. Detection sensitivity is estimated at >75%.

**Supplementary Figure 4.**
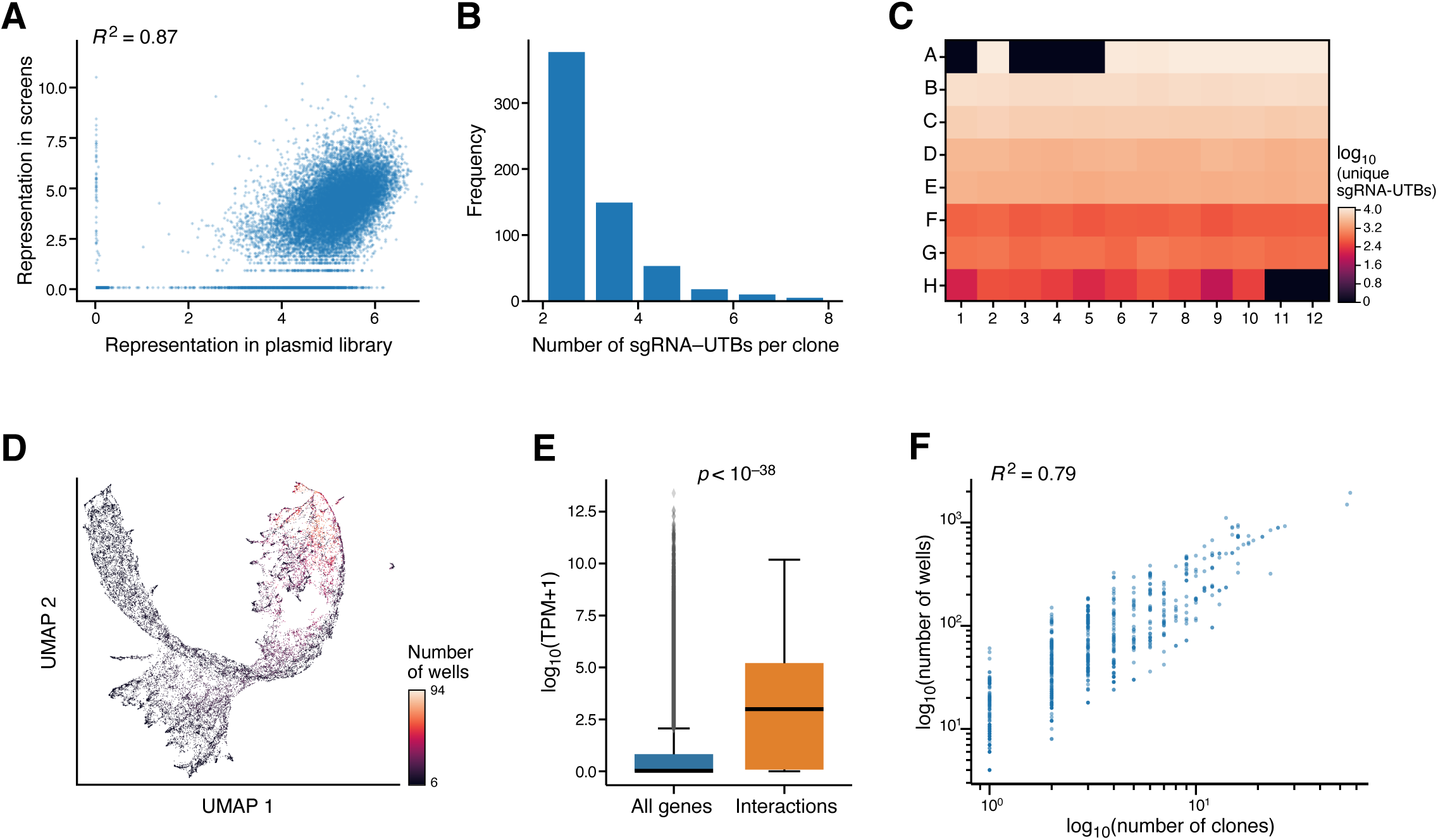
Statistics of a Shuffle-Seq screen of Vemurafenib resistance. (**a**) Library representation. Representation of genes (log_10_(number of reads)) in the initial plasmid library (*x* axis) and after transduction into cells (<u>y</u> axis). Screen representation is skewed due to limited scale of viral production in a 6-well plate (**Methods**). (**b**) Distribution of sgRNA-UTBs per clone. Number of sgRNA-UTBs observed per clone (*x* axis) in 597 clones inferred from the Shuffle-seq experiment. (**c**) sgRNA-UTBs across wells. Number of distinct sgRNA-UTBs (log_10_(number unique sgRNA-UTBs)) in each well of a 96 well plate. (**d**) Similarity structure of permuted sgRNA-UTB abundances across wells. UMAP embedding (**Methods**) of sgRNA-UTBs (dots) occurrence profiles across wells colored by number of wells in which the sgRNA-UTB is detected, after randomly permuting the sgRNA-UTB abundance matrix. (**E**) Genes involved in interactions are more likely to be expressed in A375 cells. Distribution (box: quartiles; whiskers: 1.5 interquartile range) of expression levels (*y* axis, log_10_(TPM+1)) of all genes (blue) and of the most common genes occurring in clonally inferred sgRNA-UTB pairs that survived vemurafenib selection. (**f**) The relation between the log_10_(number of clones) (*x* axis) and log_10_(number of wells) (*y* axis) in which each pair of significantly co-associated sgRNA-UTBs is present.

**Supplementary Figure Legends**

**Supplementary Table 1. Guide for the Miserly**

Spreadsheet comparing the approximate current costs associated with a standard CRISPR screen, a Shuffle-seq screen, and a single cell RNA-seq based Perturb-seq screen for a desired number of clones to be tested.

**Supplementary Table 2. sgRNA Table**

Spreadsheet listing the sgRNA oligo pools used in this paper.

## Methods

### EXPERIMENTAL PROCEDURES

#### Plasmid generation

The CROP-seq plasmid^1^ (Addgene 86708) was digested (SnaBI and NsiI) and replaced with a gBlock, which removed the sgRNA scaffold. Three plasmids containing different resistance markers (puromycin, hygromycin, and blasticidin, **Supplementary Fig. 1a**) were created from this modified CROP-seq vector. An oligonucleotide array (Custom Array, Inc.) was constructed to contain 41,467 CRISPR/Cas9 sgRNAs and partial optimized scaffolds targeting 18,377 genes in the human genome (**Supplementary Table 2**), (2.25 guides/gene). The library was depleted for sgRNAs targeting essential genes (containing only 1,981 guides targeting 1,629 essential genes), and included 10,000 control guides (targeting both intergenic regions and non-targeting guides) based on sgRNAs from previous studies^2–4^.

The oligo pool was amplified first using sub-pool specific primers, and then with the following primers to add on the remaining optimized scaffold, UTB, and subsequent plasmid homology, resulting in a 220bp product.

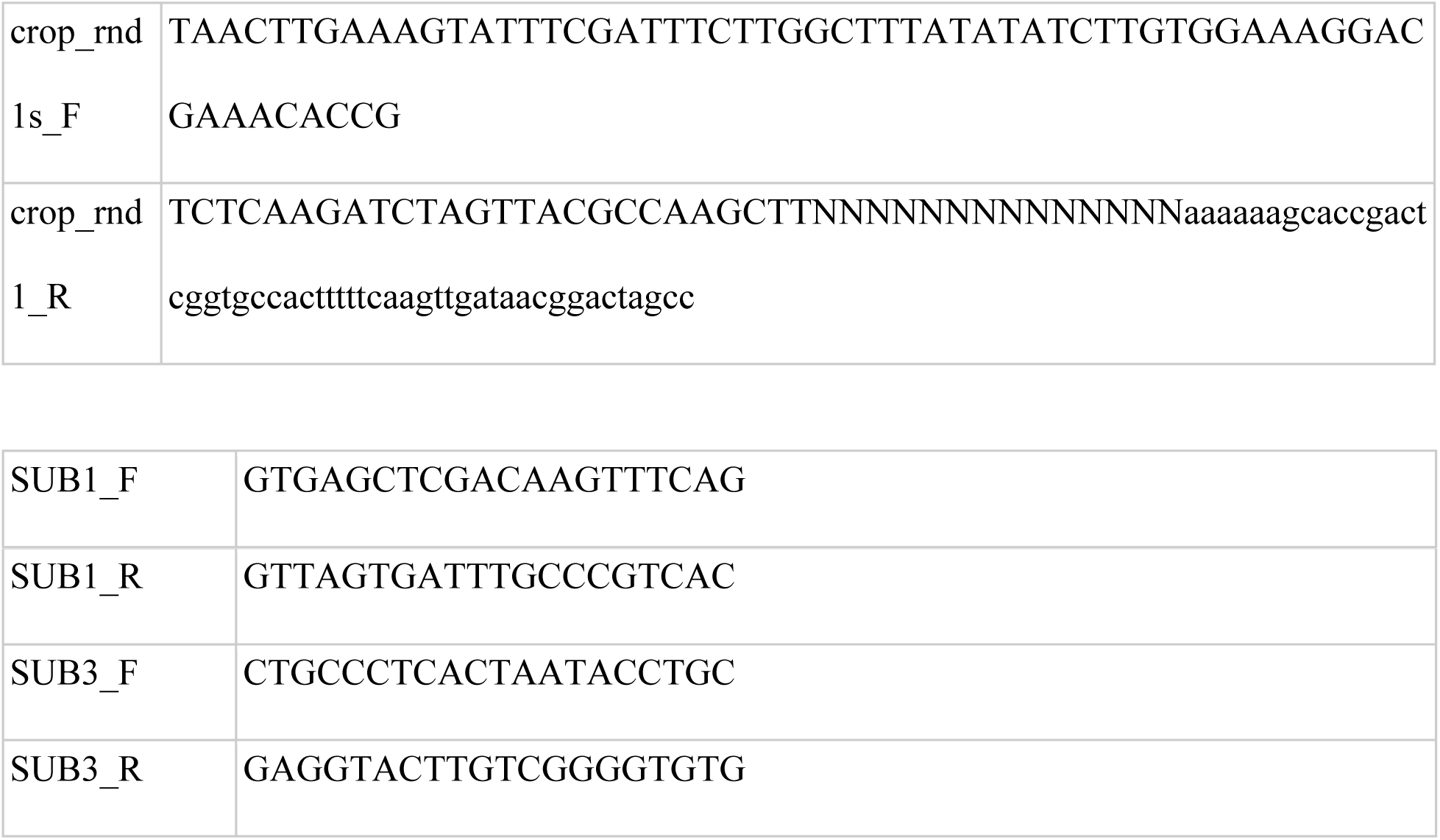

Cloning was performed as described previously.^5^

The three vectors (**Supplementary Fig.1a**) were used to generate lentiviral pools for all of our CRISPR screens as well as for the experiment validating Shuffle-seq specificity and sensitivity.

#### Lentivirus production

HEK293FT cells were seeded into 6-well plates at a density of 1 × 10^6^ cells/well and grown overnight. The following morning, the 70-80% confluent cells were transfected with 780 ng psPAX2 (Addgene plasmid #12260), 510 ng pMD2.G (Addgene plasmid #12259) and 1020 ng transfer plasmid using Lipofectamine® LTX with Plus™ Reagent (Life Technologies), according to the manufacturer’s protocol. Media were exchanged after 6 hours. Viral supernatant was harvested 24 and 48 hours following transfection, filtered through a 0.45μm polyethersulfone syringe filter (Millipore Sigma), concentrated with 100 kDA Amicon® Ultra-15 Centrifugal Filter Units (Millipore Sigma), and snap frozen at −80°C. Lentiviral stocks were titered in accordance with the alamarBlue Cell Viability Assay from the Genetic Perturbation Platform at the Broad Institute.

#### Cell culture

HEK293FT (ATCC CRL-1573) and A375 cells constitutively expressing Cas9 (a gift from the Genetic Perturbation Platform at the Broad Institute) were cultured in DMEM GlutaMAX (Thermo Fisher) supplemented with 10% heat-inactivated fetal bovine serum (Invitrogen) and 100 U/mL penicillin-streptomycin (Thermo Fisher) at 37°C, 5% CO_2_. K562 cells were cultured in RPMI 1640 Medium + GlutaMAX (Thermo Fisher) supplemented with 15% heat-inactivated FBS (Invitrogen) and 100 U/mL penicillin-streptomycin at 37°C, 5% CO_2_.

#### Estimation of the cutting efficiency of the Shuffle-seq vector

Cas9 cutting efficiency in A375 cells was measured by transducing GFP positive Cas9 containing cells with a Shuffle-seq vector targeting GFP followed by flow cytometry.

#### Validation of Shuffle-seq specificity and sensitivity

Two pools of 150,000 K562 cells without Cas9 were transduced with a pool of lentivirus expressing the Puromycin resistance gene from the Ef1α promoter in the presence of 8 µg/mL polybrene (Millipore Sigma) one at an MOI of 1 and the other at an MOI of 5. Cells were spin infected at 800 g for 45 minutes at room temperature and kept at 37°C, 5% CO_2_ overnight. The following day, the media were changed to remove the polybrene. 48 hours following transduction, cells were selected with 2 μg/mL puromycin (Thermo Fisher) for four days. Cells in the two wells were then transduced with a lentiviral pool expressing the blasticidin resistance gene from the Ef1α promoter in the presence of 8 µg/mL polybrene one at an MOI of 1 and the other at an MOI of 5 (as in the first transfection). Cells were spin infected at 800 g for 45 minutes at room temperature and kept at 37°C, 5% CO_2_ overnight. The following day, the media were changed to remove the polybrene. 48 hours following transduction, cells were selected with 6 μg/mL blasticidin for 12 days. After selection, cells from the two MOI conditions were pooled, and expanded into a T25 tissue culture flask.

As clonal controls, single cells were sorted into individual wells of one and a half 96-well plates (for a total of 144 unique clones) by flow cytometry using the Sony SH800 Cell Sorter instrument, and allowed to clonally expand for a month. After library construction, wells with low cell numbers were removed from subsequent analysis, resulting in 88 remaining wells. Libraries were prepared and sequenced as described below.

Meanwhile, the cell pool in the T25 flask (above) was expanded into a T75 flask and passaged every 2 days to maintain the cells in exponential growth phase. In this experiment, our bottleneck size was 25,000 cells. Once the single cell clones were confluent, varying number of cells from different clones were spiked into the confluent T75 flask containing the cell pool: 5 cells from each well of row A, 10 cells from each well of row B, 25 cells from each well of row C, 75 cells from each well of row D, 150 cells from each well of row E, 300 cells from each well of row F, 750 cells from each well of row G, and 1,500 cells from each well of row H. Once the contents of the T75 flask were thoroughly mixed, 10,000 cells were plated per well in a 96 well plate. 48 hours later, cells were lysed with RLT Plus buffer and stored at −80°C for downstream Shuffle-seq library construction as described below.

#### Melanoma drug resistance screen

100,000 A375 cells containing Cas9 were plated in each well of a 12 well plate. Wells were transduced with a lentiviral pool expressing the puromycin resistance gene from the Ef1α promoter in the presence of 8 µg/mL polybrene (Millipore Sigma) at a MOI of 1, 6, or 16 (4 wells per each MOI). Cells were spin infected at 1,400g for 2 hours at 33°C and kept at 37°C, 5% CO_2_ overnight. 24 hours after transduction, media were exchanged to remove polybrene. Cells were then co-selected with 1 μg/mL puromycin (Thermo Fisher) and 2 μg/mL of blasticidin (for Cas9 expression).

A week after transduction, cells were transduced with a hygromycin lentiviral pool. Each well was transduced at the same MOI as before. 24 hours after transduction, media was exchanged to remove polybrene. Cells were then selected with 300 μg/mL hygromycin (Thermo Fisher) for 10 days. Two days later, 1,875,000 cells were expanded into a T75 flask and treated with 2 μM Vemurafenib (PLX4032; Selleckchem).

On the following day, single cells were sorted into individual wells of a 96-well plate by flow cytometry using the Sony SH800 Cell Sorter instrument and allowed to clonally expand for a month in order to make an empirical estimate of MOI.

Three weeks after initiation of drug treatment, the cell density of the Vemurafenib-dosed cells in the T75 flask was measured on the hemocytometer, and the cells were manually pipetted into each well of a 96-well plate to achieve the following densities:

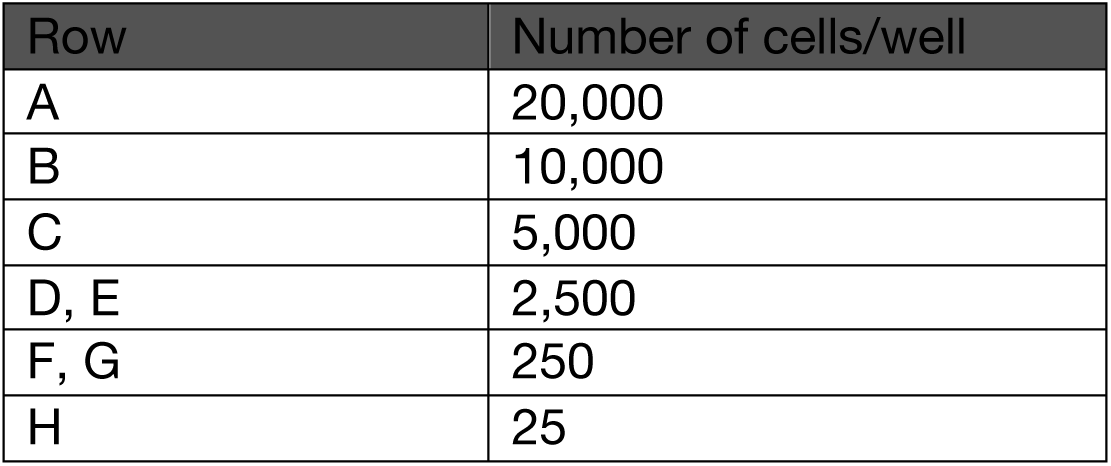

| Row | Number of cells/well |
| --- | --- |
| A | 20,000 |
| B | 10,000 |
| C | 5,000 |
| D, E | 2,500 |
| F, G | 250 |
| H | 25 |

Cells were then pelleted by centrifugation, lysed in RLT Plus buffer. Shuffle-seq library prep was performed as described below. Finally, the plate was sequenced on an Illumina Nextseq.

#### sgRNA library construction and sequencing

RNA was isolated from the cell lysate using Agencourt RNAClean XP SPRI beads (Beckman Coulter) in a 2X ratio. Each sample was eluted in a mix composed of: 3.55μL nuclease-free water, 0.1μL of 100 μM TSO, 0.5 μL of 10 mM dNTP mix (Thermo Fisher), 1 μL PEG 8000 (New England Biolabs; 50% stock), 2 μL 5X RT Buffer, 0.35 μL RNase inhibitor, 0.5 μL Maxima H Minus Reverse Transcriptase, and 2 μL RT primer. Reverse transcription was performed at 50°C for 90 minutes, followed by heat inactivation at 85°C for 5 minutes. The transcriptome was amplified by spiking in the following mix to each reaction: 12.5μL 2X KAPA HiFi HotStart ReadyMix, 0.25μL of 10μM ISPCR primer, and 2.25μL nuclease-free water. Whole Transcriptome Amplification (WTA) PCR cycling conditions were:

| Temperature | Time |  |
| --- | --- | --- |
| 98 °C | 3 min |  |
| 98 °C | 20 s | Repeat for a total of 10 cycles |
| 67 °C | 15 s |  |
| 72 °C | 6 min |  |
| 72 °C | 5 min |  |

cDNA was then purified by a 1X Agencourt AMPure XP bead clean-up (Beckman Coulter) and eluted in 15μL nuclease-free water. The yield was measured using a 96-well plate reader Qubit HS assay (Invitrogen), and the product was run on a 2% Agarose E-gel to check for the presence of a peak at 1.5-2 kb, and the absence of any fragments smaller than 500 bp.

DNA libraries were prepared by taking a fraction of the WTA product to amplify the sgRNA region and attach the Nextera adapters using the primer sequences below.

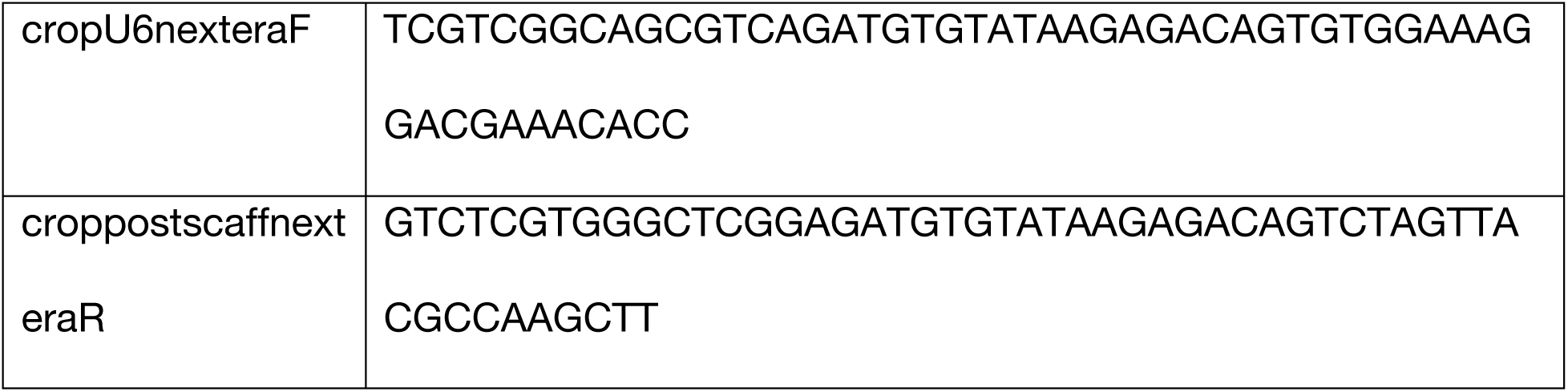

Each of the 96 PCR reactions was performed using 1-10ng of WTA, 12.5μL NEBNext® High-Fidelity 2X PCR Master Mix, 2.5μL of 10μM primer mix, and nuclease-free water (for a final volume of 25 μL). The optimal number of cycles was determined by qPCR. The PCR cycling conditions were:

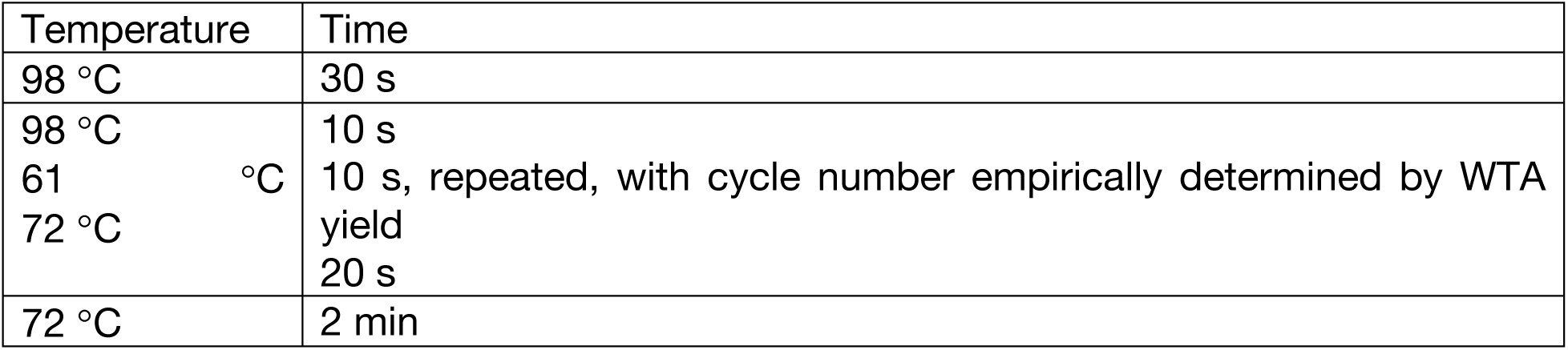

The resulting PCR 1 product was purified by a 0.8X Agencourt AMPure XP bead clean-up (Beckman Coulter). The yield was measured using a Qubit HS assay, and the size was verified by running the product on a 2% Agarose E-gel. Next, the PCR 1 product was further amplified to attach barcoded Nextera indices. Each of the 96 PCR reactions was performed using 1-10ng of PCR 1 product, 12.5 μL NEBNext® High-Fidelity 2X PCR Master Mix, 2.5μL of 10 μM Nextera P5/P7 primer mix, and nuclease-free water (for a final volume of 25μL). The optimal number of cycles was again determined by qPCR. The PCR cycling conditions were:

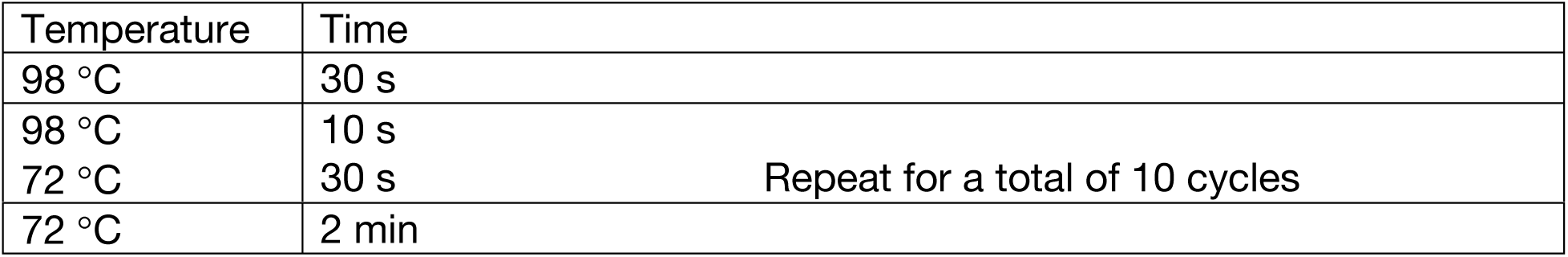

The barcoded PCR 2 products were then pooled and purified twice by a 0.8X Agencourt AMPure XP bead clean-up to remove any residual primer or primer-dimer. The yield was measured using a Qubit HS assay, and the size was verified by running the product on a 2% Agarose E-gel.

Pair-end sequencing was performed on a NextSeq™ 500 instrument. The run parameters were: 42 cycles for read 1 (sgRNA), 34 cycles for read 2 (UTB), 8 cycles for index 1, and 8 cycles for index 2. To improve run quality and base clustering, 10-50% of PhiX Control v3 Library was spiked into each run.

### COMPUTATIONAL ANALYSIS

#### Data pre-processing

Sequencing data was demultiplexed using *bcl2fastq* and a sample sheet containing the index barcodes for each well of the 96 well plate. Reads for each well were aligned using bowtie2^6^ with a reference genome consisting of a portion of the modified CROP-seq vector centered on the sgRNA-UTB construct with 60bp flanking either end. The UTB was replaced with N (ambiguous bases). After alignment, reads with fewer than two mismatches to their corresponding sgRNA were used for subsequent analysis. The UTB section was extracted from read 2 (the 14 bp downstream of the constant priming region) and counts of sgRNA-UTB were created by using Linux command uniq -c for the sgRNA-UTB pair extracted from the BAM file. This process was performed for each well and the resulting sgRNA-UTB abundance vectors were compiled by a custom Python script to create a sgRNA-UTB abundance matrix, where each column corresponded to a difference well.

#### Normalization of sgRNA-UTB matrix

After sequencing of a Shuffle-seq library, a matrix of sgRNA-UTB abundances across wells was generated. PCR chimeras (in which spurious associations between sgRNAs and UTBs are created during amplification) were removed as previously described^7^. Read depth normalization was performed by normalizing the read counts for each well to sum to 1 million (TPM). Finally, a log2 transform was performed to create a normalized sgRNA-UTB matrix.

#### Estimation of sgRNA-UTB plasmid library complexity

For a Shuffle-seq experiment, sgRNA-UTB barcode complexity should significantly exceed the number of clones in the experiment. The number of reads per sgRNA-UTB was used to estimate a zero-truncated Poisson parameter (*λ*). The log-likelihood was evaluated as:

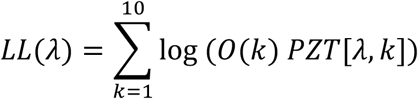

Where *O(k)* is the number of sgRNA-UTBs with *k* reads detected and *PZT*[*λ, k*] is a zero-truncated Poisson distribution. Library complexity was estimated as the total number of reads divided by the estimated *λ*.

#### Statistical inference of clones

To infer clonal associations between pairs of sgRNA-UTBs, an empirical null distribution was generated by randomly permuting (up to 100 times) the normalized sgRNA-UTB by well matrix. In an HDR-Shuffle-seq experiment (with variable numbers of cells per well), the permutation is performed only between those wells that have received similar number of cells (in the case of the vemurafenib experiment for each row of the 96 well plate). This permutation procedure ensures an empirical distribution that reflects variation in conditional distributions for the rows and columns of the sgRNA-UTB by well matrix.

Only sgRNA-UTBs that occur in similar number of wells need to be tested, thus reducing both computational complexity and the number of hypotheses tested in the family. Overlapping sets of sgRNA-UTBs whose well abundance was within a factor of 2 of one another were evaluated. For each set, pairwise Pearson correlation coefficients were calculated for each pairs of sgRNA-UTBs for the actual and permuted corresponding sgRNA-UTB submatrices, respectively. Empirical p-values for each pair of sgRNA-UTB were estimated by evaluating their real correlation as a percentile in the corresponding subset of the permuted distribution. A Benjamini-Hochberg procedure was used to control for the False Discovery Rate and a clonal pair is reported if its corresponding q-value is less than 0.05.

#### Estimation of specificity/sensitivity for 88 clones

In order to determine the sensitivity and specificity of the Shuffle-seq approach. The ability to recover sgRNA-UTBs that were known to co-associate based on 88 single cell clones was evaluated. The statistical inference procedure was performed as above, The permuted sgRNA-UTB was constructed using both all sgRNA-UTBs known not to be present in the 88 single cell clones and the sgRNA-UTBs known to be present in the 88 single cell clones. For both approaches the AUC of recovering true sgRNA-UTB pairs was >0.996.

## References

1. Tong, A. Y. et al. Global Mapping of the Yeast Genetic Interaction Network. 303, 808–814 (2004).

2. Costanzo, M. et al. The Genetic Landscape of a Cell. Science (80-.). 327, 425–431 (2010).

3. Costanzo, M. et al. A global genetic interaction network maps a wiring diagram of cellular function. Science 353, (2016).

4. Kuzmin, E. et al. Systematic analysis of complex genetic interactions. Science (80-.). 360, eaao1729 (2018).

5. Hartl, D. L. What can we learn from fitness landscapes? Curr. Opin. Microbiol. 21, 51–57 (2014).

6. Weinreich, D. M., Lan, Y., Wylie, C. S. & Heckendorn, R. B. Should evolutionary geneticists worry about higher-order epistasis? Curr. Opin. Genet. Dev. 23, 700–707 (2013).

7. Takahashi, K. & Yamanaka, S. Induction of pluripotent stem cells from mouse embryonic and adult fibroblast cultures by defined factors. Cell 126, 663–76 (2006).

8. Shen, J. P. et al. Combinatorial CRISPR – Cas9 screens for de novo mapping of genetic interactions. 14, (2017).

9. Najm, F. J. et al. Orthologous CRISPR – Cas9 enzymes for combinatorial genetic screens. Nat. Publ. Gr. 36, 179–189 (2018).

10. Han, K. et al. Synergistic drug combinations for cancer identified in a CRISPR screen for pairwise genetic interactions. (2017). doi:10.1038/nbt.3834

11. Wong, A. S. L. et al. Multiplexed barcoded CRISPR-Cas9 screening enabled by CombiGEM. Proc. Natl. Acad. Sci. 113, 2544–2549 (2016).

12. Bassik, M. C. et al. A systematic mammalian genetic interaction map reveals pathways underlying ricin susceptibility. Cell 152, 909–22 (2013).

13. Laufer, C., Fischer, B., Billmann, M., Huber, W. & Boutros, M. Mapping genetic interactions in human cancer cells with RNAi and multiparametric phenotyping. Nat. Methods 10, 427–31 (2013).

14. Dixit, A. et al. Perturb-Seq: Dissecting Molecular Circuits with Scalable Single-Cell RNA Profiling of Pooled Genetic Screens. Cell 167, 1853-1866.e17 (2016).

15. Adamson, B. et al. A Multiplexed Single-Cell CRISPR Screening Platform Enables Systematic Dissection of the Resource A Multiplexed Single-Cell CRISPR Screening Platform Enables Systematic Dissection. 1867–1882 (2016). doi:10.1016/j.cell.2016.11.048

16. Jaitin, D. A. et al. Dissecting Immune Circuits by Linking CRISPR-Pooled Screens with Single-Cell RNA-Seq Resource Dissecting Immune Circuits by Linking CRISPR-Pooled Screens with Single-Cell RNA-Seq. Cell 167, 1883-1888.e15 (2016).

17. Datlinger, P. et al. Pooled CRISPR screening with single-cell transcriptome readout. Nat.Methods 14, 297–301 (2017).

18. Kumar, M., Keller, B., Makalou, N. & Sutton, R. E. Systematic Determination of the Packaging Limit of Lentiviral Vectors. Hum. Gene Ther. 12, 1893–1905 (2001).

19. Sack, L. M., Davoli, T., Xu, Q., Li, M. Z. & Elledge, S. J. Sources of Error in Mammalian Genetic Screens. G3&#58; Genes|Genomes|Genetics 6, 2781–2790 (2016).

20. Michlits, G. et al. CRISPR-UMI: single-cell lineage tracing of pooled CRISPR–Cas9 screens. Nat. Methods 14, 1191–1197 (2017).

21. Schmierer, B. et al. CRISPR/Cas9 screening using unique molecular identifiers. Mol. Syst. Biol. 13, 945 (2017).

22. Howie, B. et al. High-throughput pairing of T cell receptor and sequences. Sci. Transl. Med. 7, 301ra131–301ra131 (2015).

23. Chen, B. et al. Dynamic Imaging of Genomic Loci in Living Human Cells by an Optimized CRISPR/Cas System. Cell 155, 1479–1491 (2013).

24. Shalem, O. et al. Genome-scale CRISPR-Cas9 knockout screening in human cells. Science 343, 84–7 (2014).

25. Morgens, D. W. et al. Genome-scale measurement of off-target activity using Cas9 toxicity in high-throughput screens. Nat. Commun. 8, 1–8 (2017).

26. Cleary, B., Cong, L., Cheung, A., Lander, E. S. & Regev, A. Efficient Generation of Transcriptomic Profiles by Random Composite Measurements. Cell 171, 1424-1436.e18 (2017).

27. Campa, C. C., Weisbach, N. R., Santinha, A. J., Incarnato, D. & Platt, R. J. Multiplexed genome engineering by Cas12a and CRISPR arrays encoded on single transcripts. Nat. Methods 16, 887–893 (2019).

## References

1. Datlinger, P. et al. Pooled CRISPR screening with single-cell transcriptome readout. Nat. Methods 14, 297–301 (2017).

2. Doench, J. G. et al. Optimized sgRNA design to maximize activity and minimize off-target effects of CRISPR-Cas9. Nat. Biotechnol. 34, 184–191 (2016).

3. Horlbeck, M. A. et al. Compact and highly active next-generation libraries for CRISPR-mediated gene repression and activation. Elife 5, 1–20 (2016).

4. Morgens, D. W. et al. Genome-scale measurement of off-target activity using Cas9 toxicity in high-throughput screens. Nat. Commun. 8, 1–8 (2017).

5. Fulco, C. P. et al. Systematic mapping of functional enhancer–promoter connections with CRISPR interference. Science (80-.). 354, 769–773 (2016).

6. Langmead, B. & Salzberg, S. L. Fast gapped-read alignment with Bowtie 2. Nat. Methods 9, 357–359 (2012).

7. Dixit, A. Correcting Chimeric Crosstalk in Single Cell RNA-seq Experiments. bioRxiv (2016). doi:10.1101/093237

